# Understanding and eliminating the detrimental effect of endogenous thiamine auxotrophy on metabolism of the oleaginous yeast *Yarrowia lipolytica*

**DOI:** 10.1101/753525

**Authors:** Caleb Walker, Seunghyun Ryu, Richard J. Giannone, Sergio Garcia, Cong T. Trinh

## Abstract

Thiamine is an essential vitamin that functions as a cofactor for key enzymes in carbon and energy metabolism for all living cells. While most plants, fungi and bacteria can synthesize thiamine *de novo*, the oleaginous yeast, *Yarrowia lipolytica*, cannot. In this study, we used proteomics together with physiological characterization to understand key metabolic processes influenced and regulated by thiamine availability and identified the genetic basis of thiamine auxotrophy in *Y. lipolytica*. Specifically, we found thiamine depletion results in decreased protein abundance of the lipid biosynthesis pathways and energy metabolism (i.e., ATP synthase), attributing to the negligible growth and poor sugar assimilation observed in our study. Using comparative genomics, we identified the missing gene scTHI13, encoding the 4-amino-5-hydroxymethyl-2-methylpyrimidine phosphate synthase for the *de novo* thiamine synthesis in *Y. lipolytica,* and discovered an exceptional promoter, P3, that exhibits strong activation or tight repression by low and high thiamine concentrations, respectively. Capitalizing on the strength of our thiamine-regulated promoter (P3) to express the missing gene, we engineered the first thiamine-prototrophic *Y. lipolytica* reported to date. By comparing this engineered strain to the wildtype, we unveiled the tight relationship linking thiamine availability to lipid biosynthesis and demonstrated enhanced lipid production with thiamine supplementation in the engineered thiamine-prototrophic *Y. lipolytica*.

**IMPORTANCE:** Thiamine plays a crucial role as an essential cofactor for enzymes in carbon and energy metabolism of all living cells. Thiamine deficiency has detrimental consequences on cellular health. *Yarrowia lipolytica*, a non-conventional oleaginous yeast with broad biotechnological applications, is a native thiamine auxotroph, whose effects on cellular metabolism are not well understood. Therefore, *Y. lipolytica* is an ideal eukaryotic host to study thiamine metabolism, especially as mammalian cells are also thiamine-auxotrophic and thiamine deficiency is implicated in several human diseases. This study elucidates the fundamentals of thiamine deficiency on cellular metabolism of *Y. lipolytica* and identifies genes and novel thiamine-regulated elements that eliminate thiamine auxotrophy in *Y. lipolytica*. Furthermore, discovery of thiamine-regulated elements enables development of thiamine biosensors with useful applications in synthetic biology and metabolic engineering.

## INTRODUCTION

Thiamine, or vitamin B1, was the first B vitamin discovered. Its activated form, thiamine pyrophosphate (TPP), functions as a cofactor for key enzymes in carbon metabolism including glycolysis, citrate cycle, pentose phosphate and branched chain amino acid pathways (Figure 1) (1). TPP-dependent enzymes, including pyruvate dehydrogenase (PDH), α-ketoglutarate dehydrogenase (KGDH), transketolase (TKL), branched chain α-ketoacid dehydrogenase (BCKDC) and acetolactate synthase (AHAS), are essential for maintaining cell growth and preventing metabolic stress (2, 3). Specifically, PDH links the glycolysis and citrate (Krebs) cycle by catalyzing the conversion of pyruvate into acetyl-CoA (4)(5). KGDH, the rate-limiting step of the citrate cycle, converts α-ketoglutarate to succinyl-CoA (6, 7). TKL is responsible for the last step of the pentose phosphate pathway where it interconverts pentose sugars to glycolysis intermediates (8). The pentose phosphate pathway is critical for production of ribose (i.e., RNA), precursor metabolites for aromatic amino acid pathways, and reducing equivalents (i.e., NADPH) necessary for maintaining redox balance and lipid synthesis. Hence, TKL activity is critical for RNA, protein, and lipid production while preventing oxidative stress (9, 10). Meanwhile, AHAS and BCKDC are responsible for the synthesis and degradation of branched chain amino acids (i.e., valine, leucine and isoleucine), respectively (11)(12).

**Figure 1.**
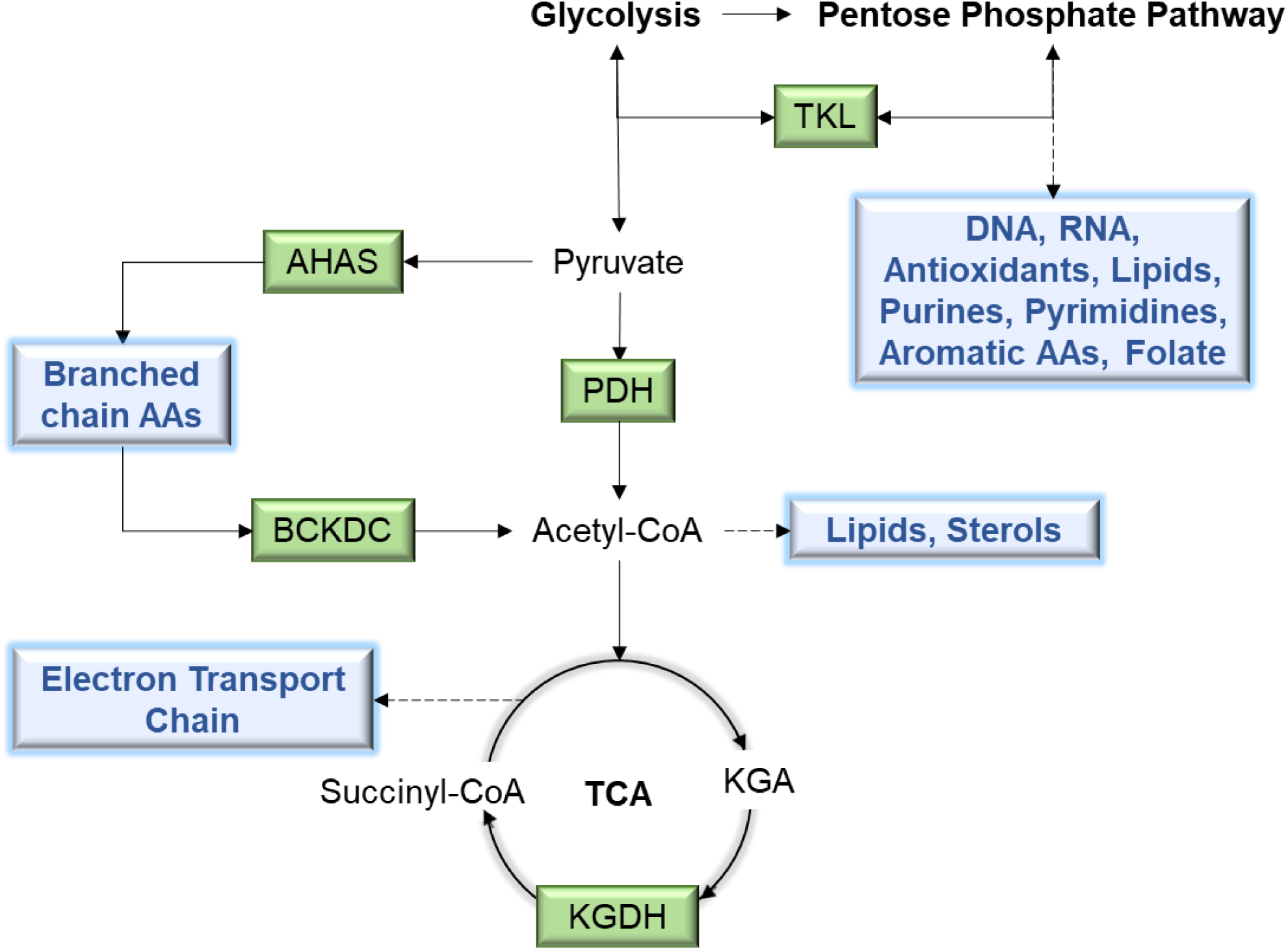
Metabolic map of thiamine-dependent enzymes (green) in relationship to central (black) and peripheral (blue) pathways. Abbreviations: TPP: thiamine pyrophosphate dependent enzymes, PDH: pyruvate dehydrogenase enzyme complex, KGDH: a-ketoglutarate dehydrogenase, TKL: transketolase, AHAS: acetolactate synthase, BCKDC: branched chain α-ketoacid dehydrogenase, TPP: thiamine pyrophosphate, AAs: amino acids, TCA: citrate (Krebs) cycle, KGA: alpha-ketoglutarate.

In mammals, thiamine deficiency affects the cardiovascular and nervous systems resulting in tremors, muscle weakness, paralysis and even death (13)(14). Thiamine deficiency can occur from inadequate intake, increased requirement, or impaired absorption of thiamine (15). Biochemical consequences of thiamine deficiency result in failure to produce adenosine triphosphate (ATP), increased acid production (i.e., lactic acid), decreased production of acetyl-compounds (i.e., acetylcholine), neurotransmitters (i.e., glutamate, aspartate, aminobutyric acid), reduced nicotinamide adenosine dinucleotide phosphate (NADH), defective ribonucleic acid (RNA) ribose synthesis, and failure to break down branched chain carboxylic acids (i.e., leucine, valine, isoleucine)(16–18).

While mammals require nutritional supplementation of thiamine from dietary sources, most bacteria, fungi and plants can synthesize thiamine endogenously. One of these exceptions is the thiamine-auxotrophic oleaginous yeast, *Yarrowia lipolytica,* which has recently emerged as an important industrial microbe with broad biotechnological applications due to its GRAS status (19), metabolic capability (20–24), and robustness (25–27). Hence, *Y. lipolytica* is an ideal eukaryotic host to study the fundamental effects of thiamine deficiency on cellular health. Currently, it is not well understood why thiamine deficiency causes failure of thiamine biosynthesis in *Y. lipolytica* and how this deficiency might directly affect other processes (e.g., lipid biosynthesis, energy metabolism, etc.) beyond what is already known.

Interestingly, *Y. lipolytica*’s auxotrophy has been exploited for enhanced production of organic acids (i.e., pyruvate and KGA) by reducing activities of PDH and KGDH under thiamine-limited conditions (28, 29). Consequently, cell growth is negatively affected by thiamine limitation and completely prevented when thiamine is depleted from the media. Not surprisingly, the genes in thiamine metabolism are tightly regulated by thiamine concentrations (30, 31). To this end, numerous thiamine-regulated promoters have been discovered in yeast that enable control of genetic expression by adjusting thiamine concentration in the culture media (32–34). However, endogenous thiamine-regulated promoters have not yet been discovered in *Y. lipolytica*. Furthermore, fundamental understanding of how thiamine deficiency inhibits metabolism and hence growth of *Y. lipolytica* is still lacking.

In this study, we shed light on the effect of thiamine deficiency on cellular metabolism in the thiamine auxotroph *Y. lipolytica*. We identified the missing gene, scTHI13, in the *de novo* thiamine biosynthesis pathway of *Y. lipolytica* and discovered a thiamine-regulated promoter, P3, that increases expression in low thiamine concentrations. By employing P3 to control the expression of scTHI13, we engineered thiamine prototrophy for the first time in *Y. lipolytica*. This enabled us to elucidate the relationship between thiamine availability and ATP and lipid biosynthesis and revealed enhanced lipid production in the engineered thiamine-prototrophic *Y. lipolytica*.

## RESULTS

### The effects of thiamine deficiency in *Y. lipolytica*

#### Cell growth and organic acid production are influenced by thiamine limitation and depletion

To demonstrate the effects of thiamine limitation, we characterized the thiamine-auxotrophic *Y. lipolytica* (YlSR001) in 0, 0.5 and 400 µg/L thiamine (Figure 2A). As expected, cell growth was inhibited by limited (0.5 ug/L thiamine) and depleted (0 ug/L thiamine) concentrations of thiamine but restored in high thiamine (400 ug/L thiamine) containing media (Figure 2B). Glucose consumption profiles were closely coordinated with cell growth (Figure 2C). Within the first 24 hr, only ∼2 g/L glucose was assimilated without thiamine while thiamine-limited cultures consumed ∼3 g/L glucose (Figure 2C). After 24 h, growth and glucose consumption were completely stalled under the no thiamine condition. In contrast, cells continued to utilize glucose even though growth was significantly inhibited.

**Figure 2.**
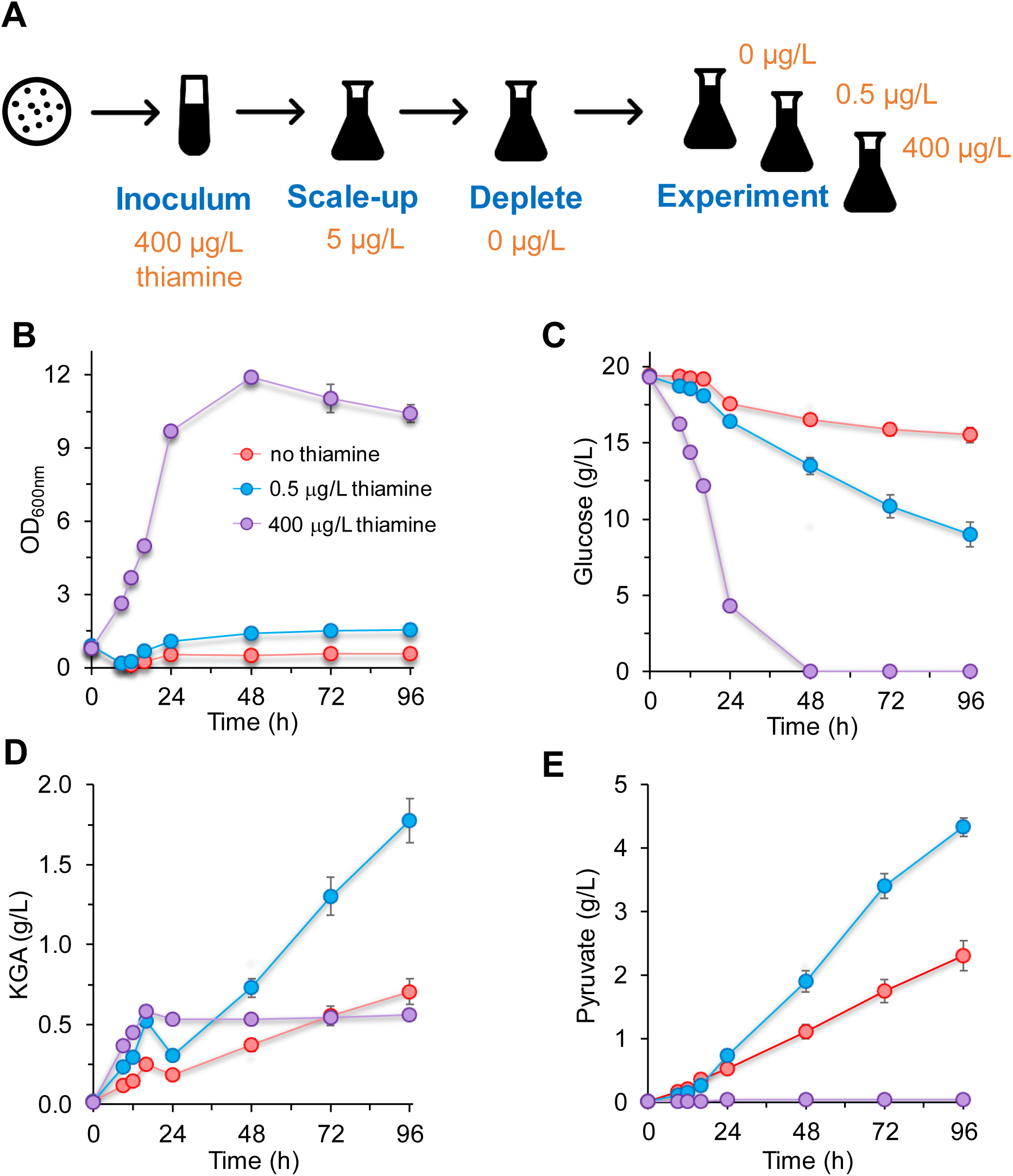
Growth characterization of the thiamine-auxotrophic *Y. lipolytica* YlSR001 in 0 (red), 0.5 (blue) and 400 (purple) µg/L thiamine. (**A**) Scheme of thiamine depletion design experiments. (**B**) Cell growth profiles. (**C**) Glucose consumption profiles. (**D**) KGA production profiles. (**E**) Pyruvate production profiles.

Next, we characterized pyruvate and KGA production since the enzymes converting these organic acids, PDH and KGDH respectively, are TPP-dependent. In high thiamine media, *Y. lipolytica* demonstrated slight KGA production (∼0.5 g/L KGA) and no pyruvate accumulation (Figure 2D, 2E). Not surprisingly, we observed substantial pyruvate titers in thiamine limited (∼4 g/L pyruvate) and depleted (∼2 g/L pyruvate) media (Figure 2E) as PDH is unable to efficiently convert pyruvate to acetyl-CoA. Similarly, KGA accumulation was enhanced in thiamine-limited media (2 g/L KGA) but KGA titers in thiamine-depleted media were comparable to high thiamine media likely due to decreased flux from glycolysis to the TCA cycle through deficient acetyl-CoA production via PDH (Figure 2D). Taken together, *Y. lipolytica* requires thiamine supplementation for cell growth and carbon assimilation but produces organic acids under thiamine-limited conditions.

#### Proteome of thiamine depletion reveals perturbation of critical metabolic pathways related to cellular growth

Next, we investigated the proteome of the thiamine-auxotrophic *Y. lipolytica* growing in 0 and 400 µg/L thiamine (Figure 3A). Across 2 exponential time points, we identified 535 upregulated and 515 downregulated proteins (i.e., log2 fold change > |1|) in response to thiamine deficiency (Figure 3B). First, we looked at metabolic enzymes that require TPP as a cofactor including PDH, KGDH, transketolase (TKL), branched chain α-ketoacid dehydrogenase (BCKDC) and acetolactate synthase (AHAS) (Supplementary Table S1). Interestingly, all proteins encoding subunits (E1-E3) of BCKDC were upregulated in thiamine depletion (Supplementary Table S1). However, none of the other TPP-requiring proteins were upregulated in media lacking thiamine except for dihydrolipoamide dehydrogenase (E3), which serves as a subunit for BCKDC, PDH and KGDH (Supplementary Table S1).

**Figure 3.**
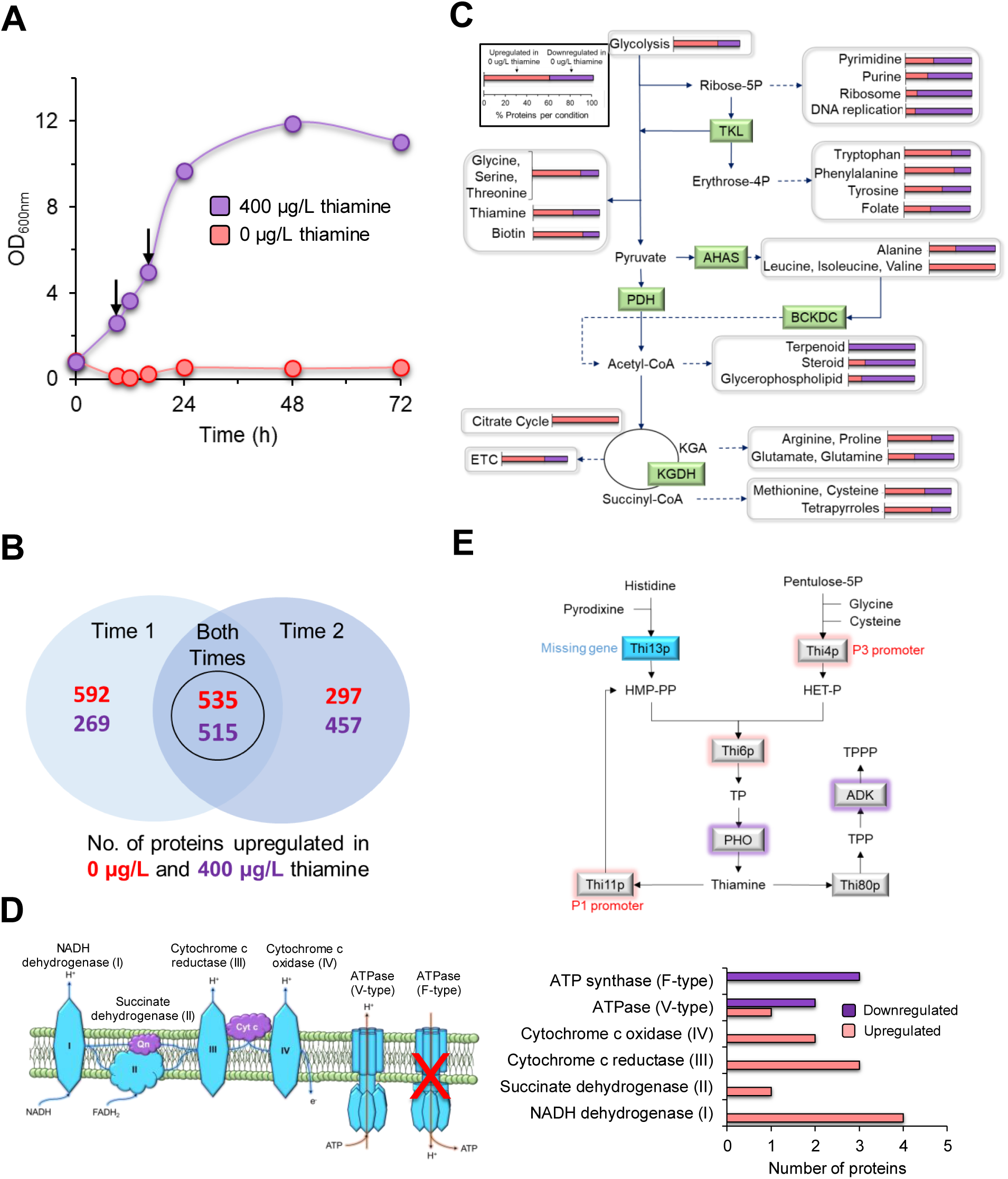
Proteomics of the thiamine-auxotrophic *Y. lipolytica* YLSR001 grown in 0 (red) and 400 (purple) µg/L thiamine. (**A**) Growth profiles and proteomics samples indicated by arrows. (**B**) Venn diagram representing upregulated and downregulated proteins at both exponential time points. (**C**) Thiamine-responsive proteomes on metabolic pathways. Bar graphs represent the percentage of proteins upregulated in 0 (red) or 400 (purple) µg/L thiamine for each pathway (**D**) Electron transport chain and ATPase. **(E**) Thiamine metabolism of *Y. lipolytica*. Locus tags: Thi4P: YALI0A09768p, Thi20p: YALI0E04224p, Thi6p: YALI0C15554p, PHO: YALI0A12573p, Thi80p: YALI0E21351p, ADK: YALI0F26521p). Abbreviations: PDH: pyruvate dehydrogenase enzyme complex, KGDH: a-ketoglutarate dehydrogenase, TKL: transketolase, AHAS: acetolactate synthase, BCKDC: branched chain α-ketoacid dehydrogenase, TPP: Thiamine pyrophosphate, HET-P: hydroxyethylthiazole phosphate, HMP-PP: hydroxymethylpyrimidine pyrophosphate, TP: thiamine monophosphate, TPP: thiamine pyrophosphate, TPPP: thiamine triphosphate.

We then mapped the 535 upregulated- and 515 downregulated proteins to their respective KEGG pathways (Figure 3C). Thiamine deficiency resulted in downregulation of proteins involved in nucleic acid metabolism (i.e., pyrimidine and purine metabolism) and genetic processes (i.e., ribosome and DNA replication). Without thiamine, cells also exhibited decreased protein abundance in lipid metabolism which includes the synthesis of terpenoids, steroids and glycerophospholipids. In contrast, thiamine deficiency resulted in upregulation of proteins contained in carbohydrate (i.e., glycolysis and citrate cycle), tetrapyrrole, biotin, aromatic (i.e., tryptophan, phenylalanine), branched chain (i.e., leucine, isoleucine, valine), and other amino acid (i.e., glycine, serine, threonine) metabolic pathways. Thiamine deficiency also affected proteins associated with the electron transport chain (ETC), amino acid (i.e., arginine, proline, glutamate, glutamine, methionine, cysteine) biosynthesis and thiamine metabolic pathways.

A closer look at proteins involved in the ETC and thiamine metabolism revealed interesting features. First, thiamine-deficient cells exhibited increased protein abundances for all four complexes of the ETC but decreased protein abundance for ATP synthase that is important for ATP generation (Figure 3D). The phenomenon correlates with the stalled assimilation of glucose that is ATP-dependent when cells grow in the absence of thiamine. Second, all but one of the proteins in thiamine metabolism were differentially regulated between thiamine-sufficient and thiamine-depleted cultures (Figure 3E). Without thiamine, we observed increased abundance of proteins in the upper branch of thiamine synthesis but decreased abundance in proteins converting thiamine monophosphate into thiamine and TPP into thiamine triphosphate. The only detected protein in thiamine metabolism that was not differentially translated was thiamine kinase (Thi90p), which is responsible for the conversion of thiamine into its activated diphosphate form, TPP. Taken together, thiamine concentration influences regulation of thiamine metabolism, whereby thiamine deficiency severely affects central carbon metabolism and elicits increased protein abundance for most carbohydrate, amino acid and energy pathways but decreased protein abundance for lipid metabolism, nucleotide metabolism and specifically, ATP synthase.

### Restoring thiamine prototrophy in *Y. lipolytica*

#### Thiamine metabolism in *Y. lipolytica* is incomplete

Next, we investigated the native thiamine metabolism of *Y. lipolytica* to elucidate underlying genetic deficiency causing thiamine-auxotrophic behavior. We compared the thiamine biosynthesis pathways between the well-characterized thiamine-prototrophic *Saccharomyces cerevisiae* and our thiamine-auxotrophic *Y. lipolytica*. Between the two organisms, only one missing gene was identified (scTHI13), encoding 4-amino-5-hydroxymethyl-2-methylpyrimidine phosphate synthase that converts histidine and pyrodixine into hydroxymethylpyrimidine pyrophosphate (HMP-PP), which is likely required for the *de novo* thiamine synthesis in *Y. lipolytica* (Figure 3E).

#### Constitutive expression of the missing thiamine gene does not effectively restore prototrophy

To enable the *de novo* biosynthesis of thiamine in *Y. lipolytica*, we constructed a vector to express scTHI13 under the constitutive promoter, TEF, frequently used for genetic overexpression in *Y. lipolytica*. Unexpectedly, expression of scTHI13 with the TEF promoter only restored the *de novo* TPP biosynthesis after days of adaptation in thiamine-depleted media (Supplementary Figure S1). Additionally, cell growth was highly deviated between replicates for both times when this experiment was conducted (Supplementary Figure S1). Though the results were somewhat promising, we endeavored to engineer a true thiamine-prototrophic *Y. lipolytica*.

#### Bioinformatics of thiamine-responsive genes reveals highly regulated thiamine promoter

We hypothesized that expression of scTHI13 was weak since the constitutive TEF promoter (35, 36) is dependent on cell growth (37, 38) and thiamine deficiency prevents cell growth. To this end, we aimed to find a promoter responsive to thiamine deficiency. Through BlastP and orthology analysis using the well-studied thiamine-regulated genes from *Pichia pastoris*, *Saccharomyces cerevisiae* and *Schizosaccharomyces pombe*, we identified three candidate genes that are putatively regulated by thiamine in *Y. lipolytica* (Supplementary Table S2). The P1 gene (YALI0E04224g) encodes a putative thiaminase that might exhibit hydroxymethylpyrimidine kinase/phosphomethylpyrimidine kinase activity for conversion of thiamine into 4-amino-5-hydroxymethyl-2-methylpyrimidine diphosphate. The P3 gene (YALI0A09768g) encodes a putative cysteine-dependent adenosine diphosphate thiazole synthase that is involved in the biosynthesis of thiazole, a thiamine precursor. Thiazole synthase converts NAD^+^ and glycine into ADP-5-ethyl-4-methylthiazole-2-carboxylate, a thiazole intermediate. Though the function of P2 (YALI0C14652g) has yet to be characterized, it harbors a NMT1 domain (Pfam:PF09084) which is required for biosynthesis of the pyrimidine moiety of thiamine (39); hence, P2 might be thiamine-regulated.

To determine if these promoters (i.e., P1, P2, and P3) are thiamine-regulated, real-time PCR was conducted to quantify mRNA levels of P1, P2, and P3 genes from YlSR001 grown in low (0.5 µg/L) and high (500 µg/L) concentrations of thiamine (Figure 4A). The expression levels of P1 and P3 genes were activated in low thiamine but inactivated in high thiamine. Remarkably, P3 exhibited substantially increased expression relative to P1 by 7.60 ± 1.51-fold in low thiamine. We did not observe transcriptional expression of P2 in low or high thiamine concentrations.

**Figure 4.**
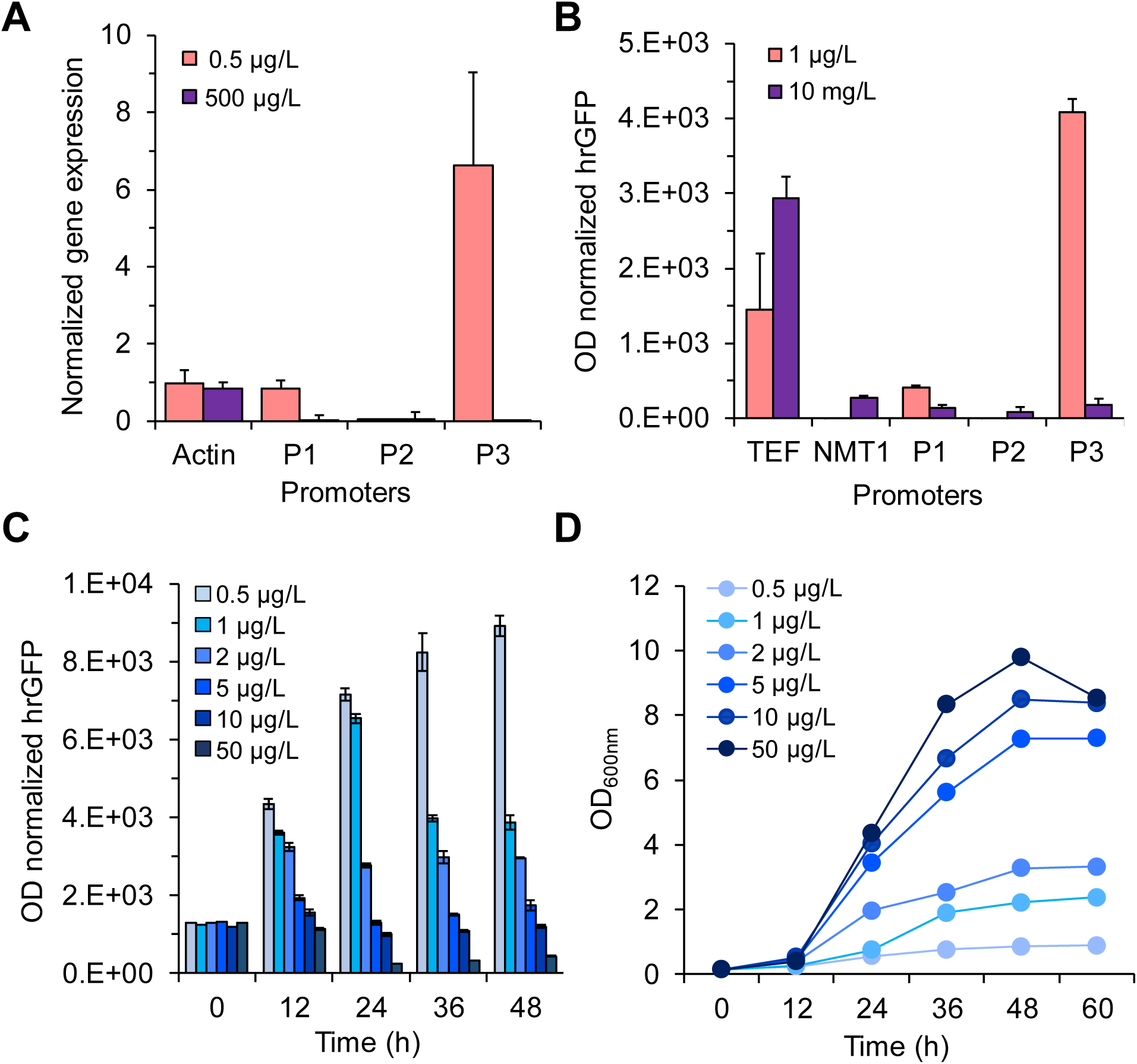
Thiamine-responsive promoter characterization. (**A**) Chromosomal gene expression of thiamine-responsive genes using rt-PCR. (**B**) hrGFP protein expression. (**C, D**) Sensitivity of thiamine-responsive promoters in presence of increasing thiamine concentrations.

We next investigated plasmid expression of these 3 putative thiamine-regulated promoters from the *Y. lipolytica* strains that harbor a humanized renilla green fluorescent protein (hrGFP)-expressing gene under the control of either P1, P2, or P3 promoter (strains YlSR1002, YlSR1003 and YlSR1004, respectively). For controls, we also built the native, constitutive TEF promoter and the heterologous NMT1 promoter from *S. pombe* that has been well-studied and applied as a thiamine-regulated promoter (40). The hrGFP intensity was monitored from cells grown at low (1 µg/L) and high (10 mg/L) concentrations of thiamine (Figure 4B). As expected, both P1 and P3 promoters were activated in low thiamine and inactivated in high thiamine. Under low thiamine conditions, the hrGFP intensity under promoter P3 was exceptionally higher than P1 and TEF promoters by 10.04 ± 0.46-fold and 2.82 ± 0.13-fold, respectively. Not surprisingly, the activity of TEF promoter was much greater in high thiamine than in low thiamine. The NMT1 promoter, however, showed no activity in low thiamine and limited activity in high thiamine.

We further investigated the sensitivity of the P3 promoter over a range of low thiamine concentrations (0.5-50 µg/L) (Figure 4C). Encouragingly, the P3 promoter exhibited tight regulation across all incremental thiamine concentrations. Of note, the activity of the P3 promoter was not affected by cell growth (Figure 4D). Taken together, these data indicate that the endogenous thiamine-regulated promoters, P1 and P3, can be applied to strain engineering efforts and are responsive to low thiamine concentrations.

#### Thiamine prototrophy is restored with thiamine regulated promoter

To restore the *de novo* thiamine synthesis in *Y. lipolytica*, we constructed the vector pSR075 to express the missing thiamine gene, scTHI13, with the thiamine-responsive promoter, P3 (i.e., YlSR1006). Remarkably, this construct demonstrated robust and reproducible growth in media lacking thiamine (Figure 5). Cell growth and glucose uptake were similar despite with low to no added thiamine (Figure 5A, 5B), but organic acid production was affected. Organic acid production in low (0.5 µg/L) and depleted (0 µg/L) thiamine conditions was diminished during the stationary phase (Figure 5C, 5D). Meanwhile, KGA accumulation only manifested in high (400 µg/L) thiamine (Figure 5C). Taken together, the thiamine prototroph strain created here, YlSR1006, grows reproducibly irrespective of thiamine concentrations but produces no organic acids in thiamine-limited media.

**Figure 5.**
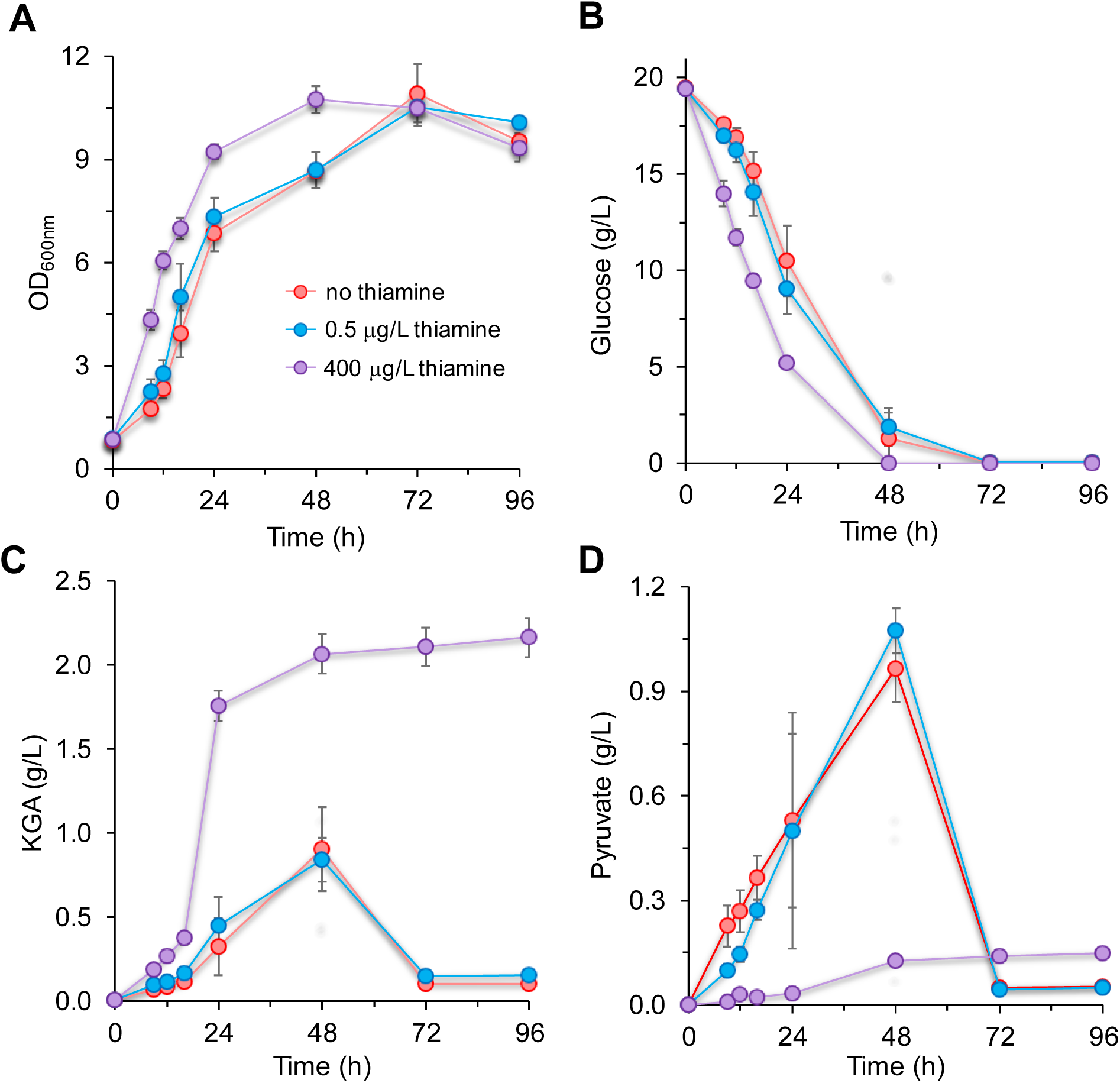
Expression of pP3-scTHI13 in YlSR1006 restores the *de novo* thiamine biosynthesis in *Y. lipolytica*. (**A**) Cell growth profiles. (**B**) Glucose consumption profiles. (**C**) KGA production profiles. (**D**) Pyruvate production profiles. Growth characterization was conducted in 0 (red), 0.5 (blue) and 400 (purple) µg/L thiamine. Abbreviations: KGA, alpha-ketoglutarate.

#### Lipid accumulation is influenced by thiamine concentrations

Finally, we investigated the relationship between thiamine availability and lipid accumulation. We cultured both thiamine-auxotrophic (wildtype) and our engineered thiamine-prototrophic (YlSR1006) strain with 0 μg/L and 400 μg/L thiamine in MpA (no nitrogen limitation, Figure 6A) and lipid production (nitrogen limitation with C:N = 100, Figure 6B) media. In non-nitrogen limited media, YlSR1006 produced more lipid than the wildtype grown with 400 μg/L thiamine supplementation (Figure 6A). Interestingly, YlSR1006 grown without thiamine accumulated lipid similar to the wildtype with 400 μg/L thiamine. However, under nitrogen-limitation, both the wildtype and YlSR1006 strains showed similar lipid production profiles when 400 μg/L thiamine was supplemented in growth medium (Figure 6B). In thiamine-lacking medium, while no lipid accumulation was expected for the wildtype, YlSR1006 was able to accumulate 3.68 ± 0.22 lipid %DCW, which was ∼50% less than YlSR1006 supplemented with 400 µg/L thiamine. Taken together, thiamine supplementation increased lipid production even for thiamine-prototroph YlSR1006, indicating thiamine plays a critical role for lipid biosynthesis in *Y. lipolytica*.

**Figure 6.**
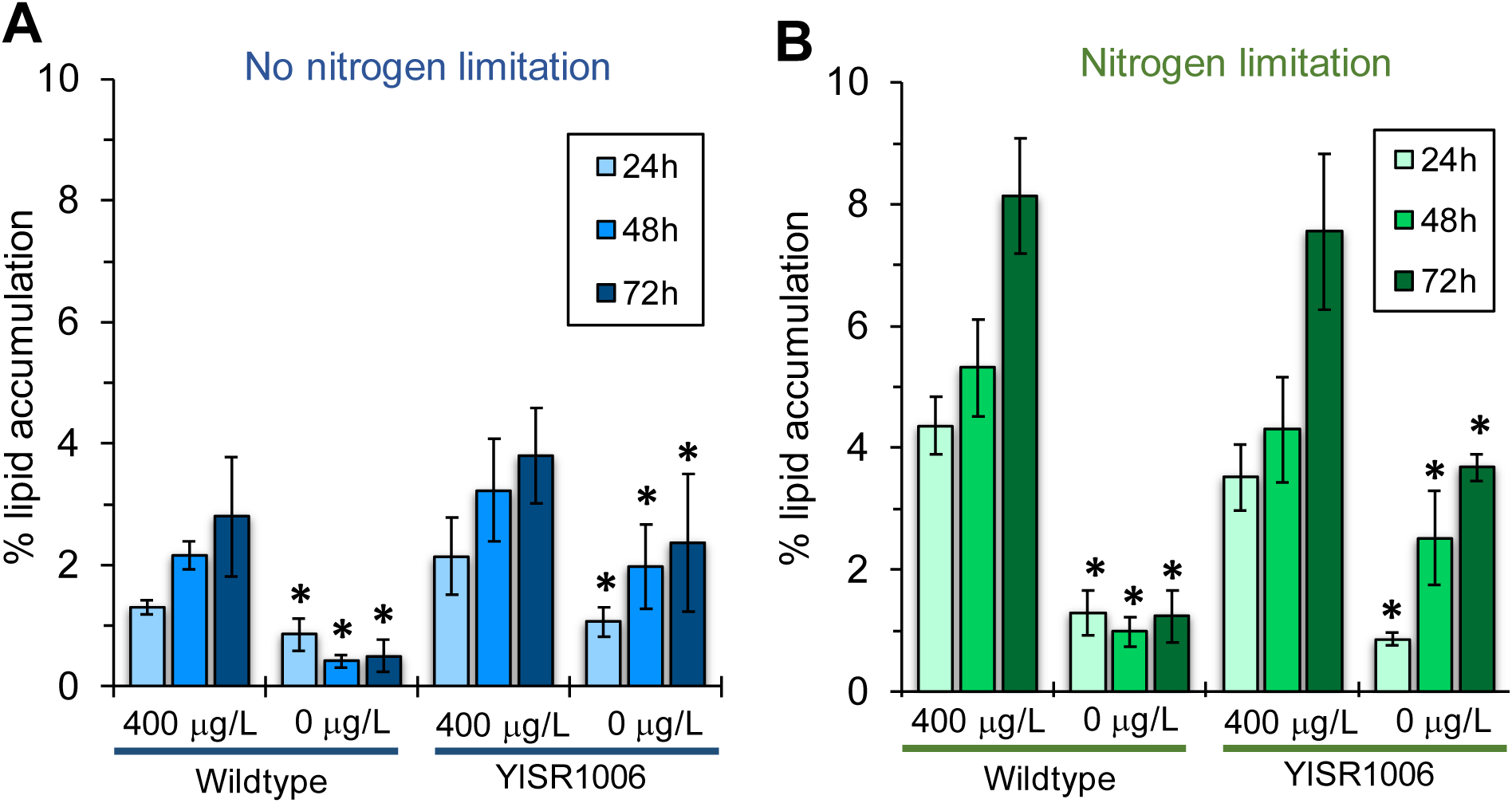
Lipid production is influenced by thiamine availability. (**A**) Lipid accumulation profiles for the thiamine-auxotrophic wildtype YlSR001 and thiamine-prototrophic strain YlSR1006 in 0 and 400 µg/L thiamine. (**B**) Lipid accumulation profiles for thiamine-auxotrophic and prototrophic strains in lipid production (C:N = 100) media with 0 and 400 µg/L thiamine. Statistical significance was calculated with one-way analysis of variance (ANOVA) with Holm-Sidak correction between 0 and 400 µg/L thiamine for each strain respectively. Symbols: “*”: p-value < 0.05.

## DISCUSSION

In this study, we observed the effects of thiamine deficiency on growth, sugar consumption, organic acid production, and proteome of the thiamine-auxotrophic *Y. lipolytica*. The activated form of thiamine, TPP, is an important cofactor for enzymes involved in vital cellular functions including energy metabolism (41), reducing oxidative and osmotic stresses (42), and catabolism of sugars (43). Hence, the consequences of thiamine deficiency are caused by the reduced activity of TPP-dependent enzymes (i.e., PDH, KGDH, TKL, AHAS and BCKDC), which ultimately leads to cell death. A comprehensive model that depicts the detrimental effect of thiamine deficiency on metabolism leading to cell death in *Y. lipolytica* is summarized in Figure 7.

**Figure 7.**
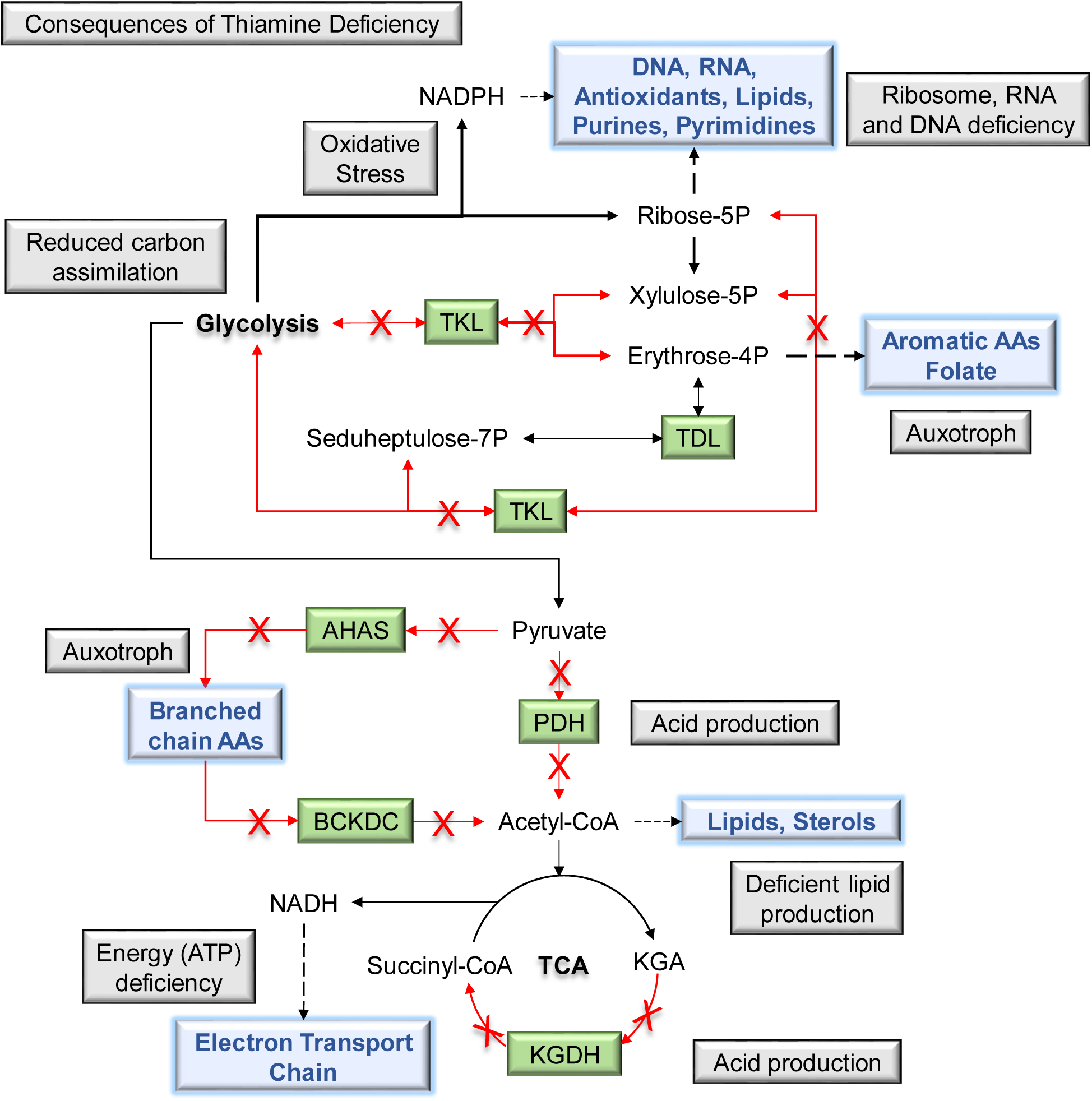
A comprehensive model explains detrimental consequences of thiamine deficiency on metabolism and cellular growth of *Y. lipolytica*.

Loss of PDH and KGDH activities result in poor growth, limited carbon assimilation and accumulation of pyruvate and KGA (Figure 2). Failure to metabolize pyruvate via PDH also inhibits production of acetyl-CoA, the precursor metabolite of the TCA cycle. The TCA cycle is further inhibited by reduced KGDH activity, preventing synthesis of NADH required for oxidative phosphorylation (i.e., ETC). Notably, in the model yeast, *Saccharomyces cerevisiae*, deletion of KGDH is known to prevent respiratory growth (44), but the exact mechanism is not well established. In our study, proteomic analysis shows that thiamine-deficient cells increased protein abundance for the first four complexes of oxidative phosphorylation (i.e., NADH dehydrogenase, succinate dehydrogenase, and cytochrome c reductase/oxidase) but decreased protein abundance for ATP synthase (Figure 3D).These new findings indicate that thiamine deficiency negatively affects respiratory energy metabolism caused by a malfunctioning TCA cycle via inhibition of PDH and KGDH as observed by inhibited growth and reduced sugar uptake in thiamine-depleted cells.

*Y. lipolytica* also decreased protein abundances in glycerophospholipid, terpenoid backbone and sterol biosynthesis pathways in thiamine-deficient cells (Figure 3C). These phenomena are likely consequences of reduced PDH activity (i.e., reduced pools of acetyl-CoA) and likely affected by NADPH production by the pentose phosphate pathway. In the pentose phosphate pathway, TKL interconverts pentose sugars and hexose sugars, the later which serve as glycolytic intermediates (e.g., fructose-6P, glyceraldehyde-3P). Hence, loss of TKL activity likely effects the production of ribose-5P and NADPH which are required for synthesis of lipids, RNA, DNA, purines, pyrimidines, and antioxidants (45). Interestingly, loss of TKL activity also prevents production of erythrose-4P, impeding synthesis of folate and aromatic amino acids (i.e., phenylalanine, tyrosine, and tryptophan) as previously observed in TKL-deletion mutants of *S. cerevisiae* (46).

In our study, *Y. lipolytica* also responded to thiamine deficiency by increasing protein abundance of enzymes involved in branched chain α-amino acid metabolism (i.e., valine, leucine and isoleucine). Consistent with the previous studies with *S. cerevisiae*, deletion of BCKDC results in branched chain α-amino acid auxotrophic phenotypes (47). Interestingly, BCKDC was the only TPP-requiring enzyme with all 3 subunits upregulated in our thiamine-depleted *Y. lipolytica* cells. However, this finding might be the result of leucine supplementation in the media since our *Y. lipolytica* strain is leucine auxotrophic. Taken together, thiamine deficiency inhibited cell growth severely limiting energy production, amino acids (i.e., aromatic and branched chain) and lipid synthesis, ultimately leading to cell death (Figure 7).

Despite thiamine-mediated growth inhibition, *Y. lipolytica* exhibited strong upregulation of proteins contained within thiamine metabolism in response to thiamine depletion (Figure 3E). Interestingly, one of the enzymes upregulated in thiamine depletion, cysteine-dependent adenosine diphosphate thiazole synthase (YALI0A09768g), is promoted by our thiamine-regulated promoter, P3. While various constitutive and inducible promoters are available for *Y. lipolytica* (48–51), our tightly regulated, thiamine-responsive P3 promoter is especially useful for strong, inducible expression in low thiamine conditions. Under optimized conditions, the activity of pP3 was 2.82 ± 0.13-fold higher than TEF (Figure 4B). Hence, gene overexpression using the P3 promoter is highly desirable over the constitutive TEF promoter in low thiamine concentrations. This was demonstrated by restoring thiamine prototrophy in *Y. lipolytica* using P3 promoter (Figure 5), while the TEF promoter was not strong enough to accomplish stable thiamine prototrophy (Supplementary Figure S1).

Surprisingly, thiamine prototroph was restored by overexpressing 1 single gene, absent from the native metabolism of *Y. lipolytica*, for the *de novo* synthesis of thiamine (Figure 3E, Supplementary Figure S1). This begs the question, why does *Y. lipolytica* lack this gene? We hypothesize that most *Y. lipolytica* strains are isolated from thiamine-rich sources (e.g., sausage, oats, plants) (52) while most popular yeasts, including *S. cerevisiae*, are isolated from sugar-rich sources (e.g., fruits, molasses, sugarcane) (52). Remarkably, we observed enhanced lipid production by thiamine supplementation in both thiamine-auxotrophic and prototrophic strains, suggesting a relationship between lipid production and thiamine availability (Figure 6A, 6B). This relationship shows promise for industrial applications of neutral lipid production and establishes a novel direction for increasing lipid production in *Y. lipolytica*.

## MATERIALS AND METHODS

### Plasmids and strains

The list of plasmids and strains used in this study is available in Supplementary Table S3. Plasmid pSR005 carrying a humanized renilla green fluorescent protein (hrGFP) was constructed by Gibson assembly method (53) with hrGFP and pSL16-CEN1-1-227 (54). The hrGFP gene was amplified using the primers hrGFP_Fwd and hrGFP_Rev from pBABE GFP (Addgene plasmid #10668). The backbone pSL16-CEN1-1-227 was amplified with the primers pSL16_Fwd and pSL16_Rev.

Next, various promoters, TEF(404) (50), NMT1 (55), P1(1000), P2(1000), and P3(1000) were inserted into pSR005 by Gibson assembly method. Each TEF(404), P1(1000), P2(1000), and P3(1000) promoter region (size of each promoter is in parenthesis) was amplified using the primers P_TEF__Fwd/P_TEF__Rev, P_P1__Fwd/P_P1__Rev, P_P2__Fwd/P_P2__Rev, and P_P3__Fwd/P_P3__Rev, respectively from the genomic DNA of *Y. lipolytica* ATCC MYA-2613. The NMT1 promoter region was amplified using the P_NMT1__Fwd/P_NMT1__Rev primers from the genomic DNA of *S. pombe* (kindly provided by Dr. Paul Dalhaimer, Department of Chemical and Biomolecular Engineering, University of Tennessee Knoxville, TN, USA). The backbone pSR005 was amplified by using the primers pSR005_Fwd/pSR005_Rev. The constructed plasmids are pAT32y, pSR068, pSR071, pSR072, and pSR073 (Supplementary Table S3).

The plasmid pSR074 was constructed by assembly of (i) THI13sce, amplified using the primers THI13_sce__Fwd^1^/THI13_sce__Rev from the genomic DNA of *S. cerevisiae* and (ii) pSR008 backbone, amplified using the primers pSR008_Fwd/pSR008_Rev. The plasmid pSR075 was constructed by replacing the hrGFP gene with the THI13sce gene from pSR073. The THI13sce gene was amplified by using the primers THI13_sce__Fwd^2^/THI13_sce__Rev from the genomic DNA of *S. cerevisiae* and assembled with the P_P3_ promoter carrying backbone, amplified by using the primers pSR008_Fwd/pSR073_Rev.

The *Y. lipolytica* ATCC MYA-2613, obtained from ATCC strain collection, was used as a parent strain. YlSR101, YlSR109, YlSR1001 and-YlSR1006 (Supplementary Table S3) strains were generated by transforming the corresponding plasmids via electroporation (56). Each plasmid was transferred into *Y. lipolytica* YlSR001 via electroporation to generate strains used in this study. The YlSR109, YlSR1001-YlSR1004 strains were confirmed by the respective promoter binding forward primer along with hrGFP_Rev. The YlSR1005 and YlSR1006 strains were confirmed by TEF(−100)_Fwd or P3(−80)_Fwd together with the respective gene binding reverse primer. *Escherichia coli* TOP10 was used for molecular cloning. Primers used in this study are listed in Supplementary Table S4.

### Media and culturing conditions

#### Media

For *E. coli* culture, Luria Bertani medium containing 5 g/L yeast extract, 10 g/L tryptone, and 5 g/L NaCl, 100 mg/L ampicillin as a selection was used. For *Y. lipolytica* characterization, MpA defined media was used for all experiments. MpA components are as follows: 5 g/L (NH4)SO4, 2 g/L KH2PO4, 0.5 g/L MgSO4, 44 mg/L ZnSO4·7H2O, 79 mg/L CaCL2·2H2O, 0.8 mg/L Biotin, 100mM HEPES buffer, 100mM phosphate buffer (prepared as 0.9M dibasic, 0.1M monobasic), trace elements (0.4 mg/L ZnSO4·7H2O, 0.04 CuSO4·5H2O, 0.4 MnSO4·6H2O, 0.2 mg/L Na2MoO4·2H2O, 0.1 mg/L KI, 0.5 mg/L FeSO4, 0.5 mg/L H3BO3), 380 mg/L leucine, 20 g/L glucose and various concentrations of thiamine hydrochloride. Initial media pH was adjusted to 5pH.

#### Culturing

All experiments were conducted in a Kuhner LT-X incubator set to 28°C and 250 rpm unless otherwise stated. Fresh colonies were inoculated in 2mL of MpA medium containing 400 µg/L thiamine in 15mL culture tubes overnight. Cultures were centrifuged and resuspended in 2mL of water before transferring 1mL of this suspension into 100mL of MpA containing 5 µg/L thiamine to scale-up cultures for 2 days (Figure 2A). Next, cells were washed once with water and resuspended in 100mL of MpA lacking thiamine for 1 day to eliminate thiamine carry-over (Figure 2A). Finally, cells were washed twice with water before characterization experiments. All experiments were conducted in technical triplicates using 500mL baffled flasks unless otherwise stated.

### Analytical methods

#### Real-time quantitative PCR (rt-PCR)

*Y. lipolytica*, grown in MpA medium using glucose as a carbon source together with either low (0.5 µg/L) and high (500 µg/L) thiamine, was collected at mid-exponential phase (OD 2 ∼ 3). Total RNA was purified by using the Qiagen RNeasy mini kit (Cat # 74104, Qiagen Inc, CA, USA), and cDNA was subsequently synthesized by the QuantiTect Reverse Transcription kit (Cat # 205311, Qiagen Inc, CA, USA). To quantify mRNA expression level of genes (e.g., actin (YALI0D08272g), P1, P2, and P3), rt-PCR run was performed using the QuantiTect SYBR Green PCR kit (Cat # 204143, Qiagen Inc, CA, USA) and StepOnePlus™ Real-Time PCR System (Applied Biosystems, CA, USA). Primers used for rt-PCR are listed in Supplementary Table S4. The gene expression level was investigated after normalizing by the actin house-keeping gene as described elsewhere (23).

#### Promoter characterization with hrGFP

Fresh colonies of *Y. lipolytica* promoter constructs were grown in 2 mL of MpA medium containing 400 µg/L thiamine overnight. Cultures were washed once with water and transferred into 25mL of MpA containing 5 µg/L thiamine overnight. Finally, cells were washed twice with water before being inoculated in MpA medium with various concentrations of thiamine. Incubation was performed at 400 rpm and 28° using 96-well plates and Duetz-system covers (Cat# SMCR1296, Kuhner, Switzerland). Sacrificial samples were collected for fluorescence measurements (excitation at 485 nm and emission at 528 nm) using a synergy HT microplate reader.

#### High performance liquid chromatography

Prior to HPLC run, 1 mL of culture medium was filtered using 0.2 μm filters. Metabolites, substrates and products were quantified by a Shimadzu HPLC system equipped with UV and RID detectors (Shimadzu Scientific Instruments, Inc., MD, USA) and the Aminex 87H column (Biorad, CA, USA) with 10 mN H_2_SO_4_ mobile phase at 0.6 mL/min flow rate. The column was maintained at 48 °C (23).

#### Proteomic analysis

*Y. lipolytica* were grown in biological triplicate in 0 and 400 µg/L thiamine. Samples were collected at two time points during the exponential growth phase and processed for LC-MS/MS analysis. Whole-cell lysates were prepared by bead beating in sodium deoxycholate lysis buffer (4% SDC, 100 mM ammonium bicarbonate, pH 8.0) using 0.15 mM zirconium oxide beads and cell debris cleared by centrifugation (21,000 x g for 10 min). Crude protein concentrations were measured by a NanoDrop OneC (ThermoScientific) using absorbance at 205 nm. Samples were then adjusted to 10 mM dithiothreitol and incubated at 85 °C for 10 min to denature and reduce proteins. Cysteines were alkylated/blocked with 30 mM iodoacetamide followed by 20 min incubation at room temperature in the dark. Proteins (300 µg) were then transferred to a 10-kDa MWCO spin filter (Vivaspin 500, Sartorius) and digested *in situ* with proteomics-grade trypsin (Pierce) as previously described (57). The tryptic peptide solution was then filtered through the MWCO membrane by centrifugation (12,000 x g for 15 min), adjusted to 1% formic acid to precipitate SDC, and SDC precipitate removed from the peptide solution with water-saturated ethyl acetate. Peptide samples were then concentrated to dryness via SpeedVac, resolubilized in solvent A (5% acetonitrile, 95% water, 0.1% formic acid), and measured by NanoDrop OneC A205 to assess tryptic peptide recovery.

Peptide samples were analyzed by automated 1D LC-MS/MS analysis using a Vanquish UHPLC plumbed directly in-line with a Q Exactive Plus mass spectrometer (Thermo Scientific) outfitted with a trapping column coupled to an in-house pulled nanospray emitter. Both the trapping column (100 µm ID) and nanospray emitter (75 µm ID) were packed with 5 µm Kinetex C18 RP resin (Phenomenex) to 10 cm and 30 cm, respectively. For each sample, 3  µg of peptides were loaded, desalted, separated and analyzed across a 210 min organic gradient with the following parameters: sample injection followed by 100% solvent A chase from 0-30 min (load and desalt), linear gradient from 0% to 25% solvent B (70% acetonitrile, 30% water, 0.1% formic acid) from 30-240 min (separation), followed by a ramp to 75% solvent B from 240-250 min (wash), re-equilibration to 100% solvent A from 250-260 min and a hold at 100% solvent A from 260-280 min. Eluting peptides were measured and sequenced by data-dependent acquisition on the Q Exactive MS as previously described (57).

MS/MS spectra were searched against the *Y. lipolytica* proteome concatenated with common protein contaminants using Proteome Discover v.2.2 (ThermoScientific) employing the CharmeRT workflow (58, 59). Peptide spectrum matches (PSM) were required to be fully tryptic with 2 miscleavages; a static modification of 57.0214 Da on cysteine (carbamidomethylated) and a dynamic modification of 15.9949 Da on methionine (oxidized) residues. False-discovery rates, as assessed by matches to decoy sequences, were initially controlled at < 1% at both the PSM- and peptide-levels. FDR-controlled peptides were then quantified by chromatographic area-under-the-curve (AUC), mapped to their respective proteins, and areas summed to estimate protein-level abundance. Protein abundance distributions were then normalized across samples using InfernoRDN (60) and missing values imputed to simulate the MS instrument’s limit of detection using Perseus (61). Significant differences in protein abundance were calculated separately for each time point according to the following equation:

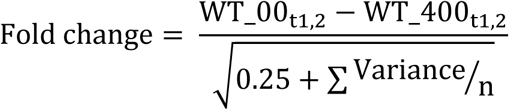

Here, WT_00 and WT_400 represent log2 normalized abundance of a protein in 0 and 400 µg/L thiamine, respectively. The denominator was used to account for error between replicates. Variance represents the variance of protein abundance between replicates, n represents the number of replicates, and 0.25 is the pseudo variance term (62). Proteins with fold changes > |1| were classified as upregulated or downregulated. Pathway annotations were performed with ClueGo (63) using KEGG (https://www.genome.jp/kegg/) database on proteins that were upregulated or downregulated at both time points.

#### Bioinformatics

Putative native *Y. lipolytica* thiamine-regulated promoters were identified by BlastP (64) and orthologs search through the KEGG sequence similarity database (https://www.kegg.jp/kegg/ssdb/). Reference genes used in this study were (i) *P. pastoris* hydroxymethylpyrimidine phosphate synthase *thi11* (PAS_chr4_0065) (65), (ii) *S. pombe* 4-amino-5-hydroxymethyl-2-methylpyrimidine phosphate synthase *nmt1* (NP_588347.1) (55), iii) *S. pombe* thiamine thiazole synthase *nmt2* (NP_596642.1) (66), and iv) *S. cerevisiae* thiamine thiazole synthase *thi4* (NP_011660.1) (67).

#### Lipid quantification

Thiamine-auxotrophic and thiamine-prototrophic strains were cultured in 0 and 400 ug/L thiamine in MpA media in triplicates as outlined previously. Lipid samples were taken from 100 uL samples of culture broth and incubated at room temperature for 15 minutes in the dark after addition of 2 uL of 1 ug/mL BODIPY (cat # D3922, Fisher Scientific) (68). Lipid standards were created by dissolving 100 mg of corn oil in 20mL ethanol and diluted from 1-.1 mg/mL prior to BODIPY staining procedure. Lipids were measured using fluorescence (ex: 485 nm/em: 528 nm) and quantified from corn oil standards. For dry cell weight (DCW) measurement, 1mL of culture broth was sampled from each replicate at each time point. Samples were centrifuged at maximum speed for 3 minutes and supernatant was discarded prior to drying samples at 55°C overnight. DCW was calculated by subtracting the dried cell pellet and tube weight by the empty tube weight. Finally, % lipid accumulation was calculated by dividing the measured lipid mg/mL by DCW mg/mL. Statistical significance was calculated using SigmaPlot 14 with one-way analysis of variance (ANOVA) with Holm-Sidak correction between 0 and 400 µg/L thiamine for each strain respectively. Symbols: “*”: p-value < 0.05.

## ACKNOWLEDGEMENTS

We would like to acknowledge financial support from the DOE BER Genomic Science Program (DE-SC0019412) and the National Science Foundation (NSF #1511881). The views, opinions, and/or findings contained in this article are those of the authors and should not be interpreted as representing the official views or policies, either expressed or implied, of the funding agencies.

