## Supplementary Materials for "Understanding and eliminating the detrimental effect of endogenous thiamine auxotrophy on metabolism of the oleaginous yeast *Yarrowia lipolytica*"

**Supplementary Table S1.** Protein expression of thiamine-dependent enzymes at both exponential-growth time points.

| Pathway | Enzyme | Subunit | Locus tag | Fold change (time 1) | Fold change (time 2) |
|---|---|---|---|---|---|
| Glycolysis | PDH | E1 | YALI0E27005g | -0.21 | -0.43 |
| | | E1 | YALI0F20702g | 0.02 | -0.33 |
| | | E2 | YALI0D23683g | -0.20 | -0.13 |
| | | E3 | YALI0D20768g | 1.62 | 2.34 |
| Citrate cycle | KGDH | E1 | YALI0E33517g | 0.92 | -0.64 |
| | | E2 | YALI0E16929g | 0.88 | -2.26 |
| | | E3 | YALI0D20768g | 1.6 | 2.34 |
| PPP | TKL | N/A | YALI0E06479g | 0.82 | -0.34 |
| Leucine, isoleucine and valine metabolism | BCKDC | E1-alpha | YALI0D08690g | 2.84 | 1.58 |
| | | E1-beta | YALI0F05038g | 2.81 | 0.79 |
| | | E2 | YALI0D23815g | 3.45 | 2.05 |
| | | E3 | YALI0D20768g | 1.62 | 2.34 |
| | AHAS | small | YALI0C09636g | -0.63 | 0 |
| | | large | YALI0C00253g | -0.31 | -0.89 |

**Supplementary Table S2.** List of putative thiamine regulatory genes in *Y. lipolytica*. Abbreviations: PP: *P. pastoris*, SC: *S. cerevisiae*, SP: *S. pombe*.

| **Genes** | **Functions** | ***Orthologs*** |
|---|---|---|
| $thi11_{PP}$ | 4-amino-5-hydroxymethyl-2-methylpyrimidine phosphate synthase | YALI0E04224g (P1) |
| $nmt1_{SP}$ | 4-amino-5-hydroxymethyl-2-methylpyrimidine phosphate synthase | YALI0C14652g (P2) |
| $nmt2_{SP}$ | thiamine thiazole synthase | YALI0A09768g (P3) |
| $thi4_{SC}$ | thiamine thiazole synthase | YALI0A09768g (P3) |

**Supplementary Table S3.** List of plasmids and strains.

| Plasmids/strains | Description | Source |
|---|---|---|
| *Plasmids* | | |
| pSL16-CEN1-1-227 | pSL16-CEN1-1-227 | (54) |
| pSR001 | pSL16-$P_{TEF}$-$T_{CYC1}$::leu2 | (26) |
| pSR008 | pSL16-$P_{TEF}$-$T_{CYC1}$::ura3 | (23) |
| pSR005 | pSL16-hrGFP-$T_{CYC1}$::leu2 | this study |
| pAT32y | pSL16-$P_{TEF}$-hrGFP-$T_{CYC1}$::leu2 | this study |
| pSR068 | pSL16-$P_{NMT1}$-hrGFP::leu2 | this study |
| pSR071 | pSL16-$P_{P1}$-hrGFP::leu2 | this study |
| pSR072 | pSL16-$P_{P2}$-hrGFP::leu2 | this study |
| pSR073 | pSL16-$P_{P3}$-hrGFP::leu2 | this study |
| pSR074 | pSL16-$P_{TEF}$-THI13sce-$T_{CYC1}$::ura3 | this study |
| pSR075 | pSL16-$P_{P3}$-THI13sce-$T_{CYC1}$::ura3 | this study |
| *Yeast strains* | | |
| YlSR001 | MATA ura3-302 leu2-270 xpr2-322 axp2-deltaNU49 XPR2::SUC2 | ATCC MYA-2613 |
| YlSR101 | YlSR001 + pSR001 | (26) |
| YlSR108 | YlSR001 + pSR008 | (23) |
| YlSR109 | YlSR001 + pAT32y | this study |
| YlSR1001 | YlSR001 + pSR068 | this study |
| YlSR1002 | YlSR001 + pSR071 | this study |
| YlSR1003 | YlSR001 + pSR072 | this study |
| YlSR1004 | YlSR001 + pSR073 | this study |
| YlSR1005 | YlSR001 + pSR074 | this study |
| YlSR1006 | YlSR001 + pSR075 | this study |

**Supplementary Table S4.** List of primers. Bold underline is AvrII sequence.

| Genes | Primers | Sequences |
|---|---|---|
| *Primers for plasmid construction* | | |
| *pSL16* | pSL16_Fwd | GCCTGCACGAGTGGGTGTAATCATGTAATTAGTTATGTCACGCTTAC |
| | pSL16_Rev | CAGGATCTGCTTGCTCACCATAGATCTGTTCGGAAATCAACGG |
| *hrGFP* | hrGFP_Fwd | AATCGGTTGAGCATCCGTTGATTTCCGAACAGATCTATGGTGAGCAAGCAGATCCTG |
| | hrGFP_Rev | GTAAGCGTGACATAACTAATTACATGATTACACCCACTCGTGCAGG |
| $P_{TEF}$ | $P_{TEF}$_Fwd | CATCCGTTGATTTCCGAACAGATCTAGAGACCGGGTTGGCGGCGTATTTG |
| | $P_{TEF}$_Rev | TTCAGGATCTGCTTGCTCACCATTTTGAATGATTCTTATACTCAGAAGGAAATGCTTAAC |
| $P_{NMT1}$ | $P_{NMT1}$_Fwd | GCATCCGTTGATTTCCGAACAGATCTTTGTATTTCAAAGGACATAATCTAAAATAATAAC |
| | $P_{NMT1}$_Rev | GGTGTTCTTCAGGATCTGCTTGCTCACCATGATTTAACAAAGCGACTATAAGTCAGAAAG |
| $P_{P1}$ | $P_{P1}$_Fwd | GGTTGAGCATCCGTTGATTTCCGAACAGATCTTGAAGTGGGTGAGTCGCCAATTATTC |
| | $P_{P1}$_Rev | CAGGCCGGTGTTCTTCAGGATCTGCTTGCTCACCATGATCGAATTGAGTCAGCGACG |
| $P_{P2}$ | $P_{P2}$_Fwd | CGGTTGAGCATCCGTTGATTTCCGAACAGATCTCAGGTGGTAGCAGCCCAAGACAATG |
| | $P_{P2}$_Rev | GGTGTTCTTCAGGATCTGCTTGCTCACCATGAATTGACGAACAGGTGTTTTGATG |
| $P_{P3}$ | $P_{P3}$_Fwd | CGGTTGAGCATCCGTTGATTTCCGAACAGATCTGAGGGGTAGTCGTAAGTTTCATC |
| | $P_{P3}$_Rev | CGGTGTTCTTCAGGATCTGCTTGCTCACCATGTTAATTGTAGGTGATATAAGGGGAAG |
| *pSR005* | pSR005_Fwd | ATGGTGAGCAAGCAGATCCTG |
| | pSR005_Rev | AGATCTGTTCGGAAATCAACGGATGCTCAAC |
| THI13$_{sce}$ | THI13$_{sce}$_Fwd[1] | CATTTCCTTCTGAGTATAAGAATCATTCAAAATGTCTACAGACAAGATCACATTTTTG |
| | THI13$_{sce}$_Fwd[2] | CACCCTTCCCCTTATATCACCTACAATTAACATGTCTACAGACAAGATCACATTTTTG |
| | THI13$_{sce}$_Rev | GAATGTAAGCGTGACATAACTAATTACATGATTAAGCTGGAAGAGCCAATCTCTTG |
| pSR008 | pSR008_Fwd | TCATGTAATTAGTTATGTCACGCTTAC |
| | pSR008_Rev | TTTGAATGATTCTTATACTCAGAAG |
| pSR073 | pSR073_Rev | GTTAATTGTAGGTGATATAAGGGGAAGG |
| *Primers for checking and sequencing* | | |
| hrGFP_seq | hrGFP_Rev | CTTGCCGCAGCCCTCCATGG |
| TEF_seq | TEF(-100)_Fwd | CACCGTCCCCGAATTACCTTTC |
| P3_seq | P3(-80)_Fwd | CTGCCGTAAATCACATACTGTCGGCTG |
| *Primers for rt-PCR* | | |
| Actin | Actin rt_Fwd | TCCAGGCCGTCCTCTCCC |
| | Actin rt_Ref | GGCCAGCCATATCGAGTCGCA |
| P1 | P1 rt_Fwd | AGGACAAGGAGCCTGCCAAG |
| | P1 rt_Rev | GGAGGCAATGGCAGAGGCTA |
| P2 | P2 rt_Fwd | TATGCAATCGGCCTCACCGA |
| | P2 rt_Rev | CTTGCCCTCCAGCTGGTCTT |
| P3 | P3 rt_Fwd | GCTGGCTCCTGTGGTCTCTC |
| | P3 rt_Rev | GAACTGCTCGGCAGGCTTTC |

**Supplementary Figure S1.** Constitutive expression of missing thiamine gene does not effectively restore thiamine prototrophy in *Y. lipolytica*. Wildtype is in blue and 6 replicates of YlSR005 growth without thiamine (0 µg/L thiamine) in orange.

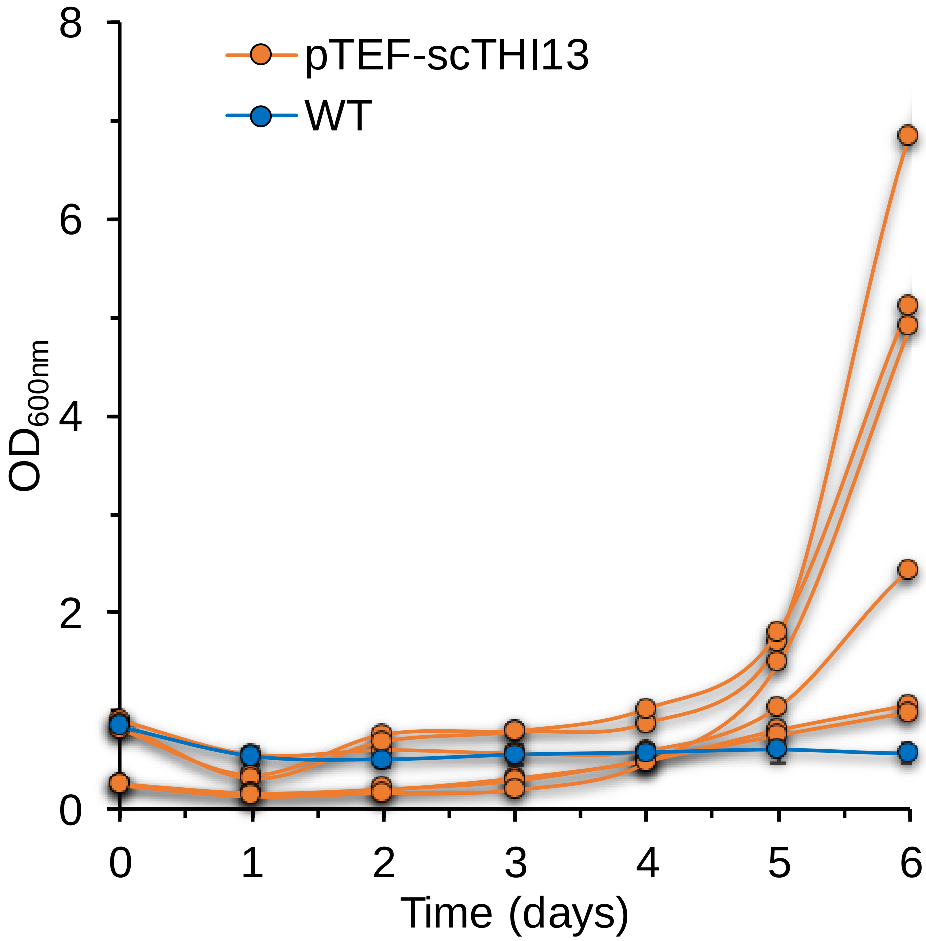
